# Model-averaged regression coefficients have a straightforward interpretation using causal conditioning

**DOI:** 10.1101/133785

**Authors:** Jeffrey A. Walker

**Affiliations:** Department of Biological Sciences, University of Southern Maine, Portland, ME 04103, USA

**Keywords:** statistical model, causal effect, probabilistic conditioning, causal conditioning, model selection

## Abstract

1. Model-averaged regression coefficients have been criticized for averaging over a set of models with parameters that have different meanings from model to model. This criticism arises because of confusion between two different parameters estimated by the coefficients of a statistical model.
2. Ever since Fisher, the textbook definition of a coefficient (a “differences in conditional means”) takes its meaning from probabilistic conditioning (*P*(*Y*|*X*)). Because the parameter estimated with probabilistic conditioning is conditional on a specific set of covariates, its meaning varies from model to model.
3. The coefficients in many applied statistical models, however, take their meaning from causal conditioning (*P*(*Y*|*do*(*X*))) and these coefficients estimate causal effect parameters (or simply, causal effects or Average Treatment Effects). Causal effect parameters are also differences in conditional expectations, but the event conditioned on is not the set of covariates in a statistical model but a hypothetical intervention. Because an effect parameter takes its meaning from causal and not probabilistic conditioning, it is the same from model to model, and an averaged coefficient has a straightforward interpretation as an estimate of a causal effect.
4. Because an effect parameter is the same from model to model, the estimates of the parameter will generally be biased. By contrast, with probabilistic conditioning, the coefficients are consistent estimates of their parameter in every model, but the parameter differs from model to model. Confounding and omitted variable bias, which are central to explanatory modeling, are meaningless in statistical modeling as mere description.
5. The argument developed here only addresses the “different parameters” criticism of model-averaged coefficients and is not advocating model averaging more generally.

## Introduction

Model averaging is an alternative to model selection (or “simplification”) for either effect estimation or prediction (Draper, 1995; Hoeting *et al.,* 1999; Burnham & Anderson, 2002). A model-averaged estimate is

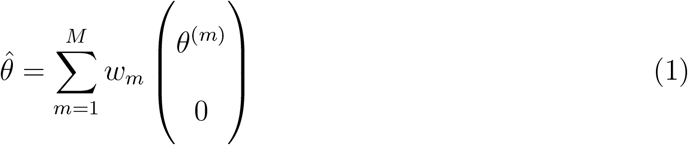

where *θ*^(*m*)^ is a parameter estimated from model *m* and *w_m_* is a weight based on some measure of model fit penalized by the number of parameters in the model. The parameter averaged could be the conditional mean (*μ_i_*) for individual *i* or the coefficient (*b_j_*) of predictor *j* (*X_j_*). Note that for coefficients, assigning *θ*^(*m*)^ = 0 when the predictor is not in the model has the effect of shrinking the estimate toward zero by an amount proportional to the weight of the model. Consequently, model averaging is an empirical shrinkage estimator and can have lower variance, at the cost of some bias, compared to the estimate from the full model (Hoeting *et al.,* 1999; Hansen, 2007). In the ecology literature, there has been some tendency to average only over models that include the parameter, in which case, the averaged coefficient is not a shrinkage estimate, but this “conditional” average is not considered further here.

Model-averaging often follows an all-subsets regression, a practice that is sometimes criticized for mindless model building. Nevertheless, averaging coefficients across subsets of the full model is a data-driven shrinkage estimator, and thus an alternative to ridge regression or Bayesian estimation, and can outperform model selection and the full model, where performance is measured by a summary of the long run frequency of error in simulations with known parameter values. Despite this apparently attractive feature of model-averaging, two recent papers in the ecology literature criticize model-averaged coefficients, not because of poor empirical performance but because the resulting coefficients are believed to have a meaningless or non-sensical interpretation. Banner & Higgs (2017) argue that the coefficients 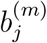 from different models estimate different parameters with different meanings and, consequently, any interpretation of an averaged coefficient is awkward at best. This “different parameter” criticism echoes the occasional comment from the statistics literature (Draper, 1999; Candolo *et al.,* 2003; Raftery & Zheng, 2003; McElreath, 2018). Cade (2015) expanded this criticism by specifically arguing that in the presence of any correlation (his “multicollinearity”) among the predictors, averaging coefficients is invalid because the units for a coefficient differ among models and, consequently, an averaged coefficient has “no defined units.” Cade’s criticism is *not* the typical caution against the estimation of regression coefficients in the presence of high collinearity that some in the literature have mistakenly believed. Instead, Cade’s criticism has only to do with the units and not with the estimation itself.

The different-parameter criticism of model-averaged regression coefficients has not been adequately addressed although there is some discussion in the statistics literature on the interpretation of the coefficients from a set of nested models. Berger *et al.* (2001) noted different potential interpretations but in the context of how to model the prior distribution and not in the context of a meaningless average. Consonni & Veronese (2008) also considered the meaning of the parameters in a sub-model and showed four different interpretations. In two of these (their interpretations 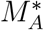 and 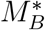), the parameter for a regression coefficient in a sub-model has the same meaning as that in the full model.

For example, if the full model is *Y_i_* = *β*_0_ + *β*_1_*X*_1*i*_ + *β*_2_*X*_2*i*_ + *ε_i_* and the submodel *Y_i_* = *β*_0_ + *β*_1_*X*_1*i*_ + *ε_i_* then *β*_1_ is the same parameter in both models if we consider the submodel to be the full model with *β*_2_ = 0. This “zero effect” interpretation is effectively the interpretation given by Hoeting *et al.* (1999) in their response to Draper (1999).

Here, I argue that model-averaged coefficients have a straightforward interpretation if the goal of the statistical model is explanation and not mere description. In short, the coefficients 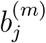 from different submodels *m* can be meaningfully averaged because they estimate a parameter common to all models. This parameter is the effect parameter, or Average Causal Effect, *β_j_*, which is the direct causal or generating effect of *X_j_* on *Y* (Morgan & Winship, 2007; Angrist & Pischke, 2008; Pearl, 2009a; Spanos, 2006b; Shalizi, 2017). While I am defending model-averaged coefficients as valid constructs with a straightforward interpretation, I am specifically not defending model-averaging as the preferred estimator of model parameters.

## Probabilistic vs. causal conditioning

A recent article on best practices involving regression-like models (Zuur & Ieno, 2016) used as an example the data of Roulin & Bersier (2007), who showed that – and entitled their paper – “nestling barn owls beg more intensely in the presence of their mother than in the presence of their father.” This title might simply be a description of the major result, that is, a difference in conditional means (on the set of covariates in the study, including time of arrival, time spent in nestbox, and a food manipulation treatment). In the discussion, however, Roulin & Bersier (2007) state that “differential begging to mother and father implies that offspring should be able to recognize the identity of each parent.” That is, the difference in chick calling is in direct response to the sex of the parent, or, put differently, the sex of the parent bringing food causally modifies the intensity of chick calling.

This example serves to introduce the major argument of the present note: The distribution of the response variable in descriptive and explanatory models are conditioned on different events and consequently, the coefficients of descriptive and explanatory models estimate different parameters (Pearl, 2009a; Shalizi, 2017; Hitchcock, 2018). In descriptive modeling, the conditional probability distribution of the response is

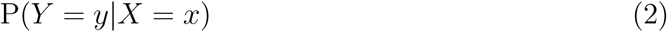

*Y* is what we observe when we also observe a specific set of *X* variables. By contrast, in explanatory modeling, the conditional probability distribution of the response is

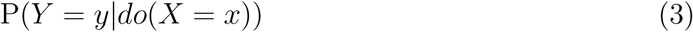

*Y* is what we would observe *after* a hypothetical intervention in the system (encoded by the *do* operator) that moved *X* to *x* (Pearl, 2009b; Shalizi, 2017; Hitchcock, 2018).

Because of this difference in conditioning, the coefficients in descriptive and explanatory models estimate different parameters. A coefficient in descriptive modeling estimates the parameter

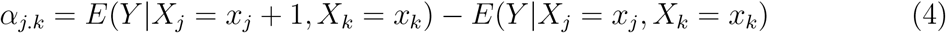

while a coefficient in explanatory modeling estimates the parameter

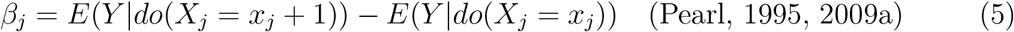

*α* is a statistical association that takes its meaning from “probabilistic conditioning” (Shalizi, 2017). *β* is a causal effect (often referred to as the “average causal effect”) that takes its meaning from “causal conditioning” (Shalizi, 2017). Or, as Judea Pearl puts it, *α* is conditioned on “seeing” while *β* is conditioned on “doing” (Pearl & Mackenzie, 2018). I refer to *α_j.k_* as a “statistical effect” or “statistical coefficient” and *β_j_* as a “causal effect” or “causal coefficient”. *α_j.k_* is often defined as a “difference in conditional means” but *β_j_* is also a difference in conditional means. Their meaning differs because the event conditioned on differs. This event is the observation of specific *X* variables for *α_j.k_* and a hypothetical intervention for *β_j_*. Importantly, the definition (or interpretation) of *β_j_* is not conditional on a set of covariates *X_k_*, even if these are measured and included in the statistical model. While I have used the *do* operator to define causal conditioning, my argument is equally valid if I used potential outcomes and the Rubin causal model (Rubin, 1974) to define causal conditioning.

While the formal recognition of probabilistic vs. causal conditioning is fairly recent, the general idea of causal conditioning goes back to the beginning of multiple regression, by George Yule, who used least squares multiple regression to estimate the causal effects of the changing demographics of pauperism of 19th century Britain (Yule, 1899; Freedman, 1999; Hepple, 2001) (partial regression coefficients were first developed as part of a “system of correlations” by Yule’s mentor and colleague Karl Pearson three years earlier). The concept of causal conditioning was expanded by the seminal work of Sewell Wright (1921, 1934) in his method of path analysis. Wright did not develop path analysis to discover causal relationships but to quantify causal effects from a pre-specified causal hypothesis in the form of paths (arrows) connecting causes (causal variables) to effects (response variables) (see below). Wright used partial regression coefficients as the effect (path) coefficients. The “difference in conditional means” interpretation of a regression coefficient began to emerge only following Fisher (1922), who seems to have been the first to conceive of regression using probabilistic conditioning (Aldrich, 2005). Pearl (2015); Pearl & Mackenzie (2018) presents a very readable history of the conflict between probabilistic and causal conditioning.

## Only causal conditioning is associated with a unique generating model

In explanatory modeling, a statistical model is the researcher’s attempt to model the true, process that generated the data. By contrast, there is no “true” generating model in probabilistic conditioning but only the specification of an idealized model from which the data could be a representative sample (Spanos, 2006a,b; Pearl, 2015). Descriptive modeling is simply not concerned with how the data were actually generated but only with the conditional probability distribution of the data once generated. To see this difference, consider a statistical model of the chick calling in nestling owls described above (simplified from the model in Zuur & Ieno (2016))

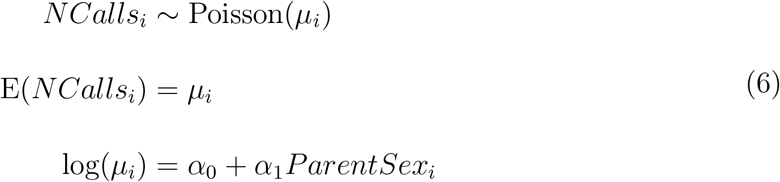

This model does not specify a generating model, only how *NCalls* is related to *ParentSex* given that we’ve observed both. It does not specify the generating model because there are, literally, an infinite number of generating models, each with a different combinations of effector (“predictor”) variables, that could generate precisely the same *P*(*NCalls*|*ParentSex*). For example, if the statistical parameters are *α* = (5.1, −0.04), then both

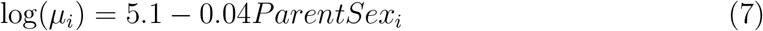

and

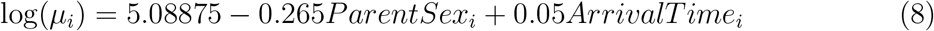

generate the same *P*(*NCalls*|*ParentSex*) (the second parameterization also requires COR(*ParentSex*, *ArrivalTime*) = 0.6, Supplement 1). In the first, the generating model (with causal parameters *β*) coincides with the statistical model. In the second, the generating model does not coincide with the statistical model. If the second model is the true generating model, and the true causal effect is −0.265, an estimate of *α*_1_ close to −0.04 is “correct” and not “wrong” if one is descriptive modeling because the goal isn’t estimation of the causal effect but the estimate of the statistical effect of *ParentSex* conditioned on no other predictors. Identifying the true set of causal effectors is not an assumption of descriptive modeling using probabilistic conditioning – that is, missing causal effectors does not violate any of the Gauss-Markov conditions.

In contrast to statistical parameters, the parameters of an explanatory model use causal conditioning and, consequently, take their meaning from causal hypotheses using substantive knowledge. Directed acyclic graphs (or path models) are an especially compact way to specify the causal hypotheses of a generating (causal) model:

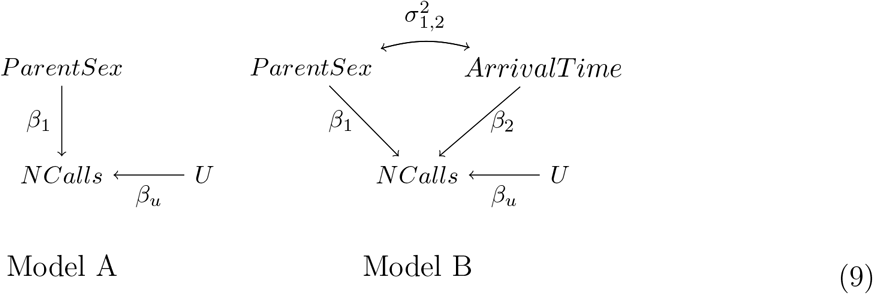

The parameters of the single-headed arrows are the (unstandardized) effect coefficients *β_j_*. The parameter of the double-headed arrow is a covariance. *U* represents all effectors that contribute to the variance in *NCalls* but are uncorrelated with the specified effectors. The meaning of *β*_1_ is the same in both model A and model B, it is the change that *would occur* in *NCalls* given an experimental intervention that sets its value from *x* to *x* + 1 (in this example, from female to male). The precise set of covariates in a statistical model used to estimate *β*_1_ has no bearing on the meaning of *β*_1_. But, in contrast to descriptive modeling, *β*_1_ is estimated without bias only when the statistical model coincides with the true, generating model. If the true generating model is Model B, and the regression model is log(*μ_i_*) = *b*_0_ + *b*_1_ *ParentSex_i_*, then the expected value of 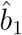 is not *β*_1_ but 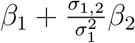.

The additional term 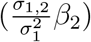 in the expectation of *b*_1_ is a bias due to the missing confounder *ArrivalTime* in the statistical model. A variable is a confounder of *X_j_* if it is both correlated with *X_j_* and has a causal effect on the response through some path that is independent of *X_j_*. In explanatory models, a modeled coefficient is a consistent estimate of the causal parameter *β_j_* only if the statistical model correctly identifies the causal structure and does not exclude confounding variables, or if the statistical model uses well-known methods to block confounders (Pearl, 2009a). By contrast, a modeled coefficient in descriptive models is not a biased estimate of *b_j.k_* if *X* with direct causal effects on *Y* are omitted from the statistical model. In fact, omitted variable bias and confounding are irrelevant or meaningless in the context of descriptive modeling, an important point that doesn’t seem to be greatly appreciated (but see, for example, Gelman & Hill, 2007; Hernán, 2018). The recognition of this distinction in ecology is unknown, but I have had reviewers of two different manuscripts (this manuscript and Walker (2014)) explain to me that omitted variable bias is not really bias because OLS is an unbiased estimator if Gauss-Markov conditions are met and omitted variables is not one of these conditions. But Gauss-Markov is not sufficient for explanatory modeling. More generally, few textbooks recognize the existence of both probabilistic and causal conditioning, and most confuse them, using something like Equation 4 to define a coefficient but then using causal vocabulary (“how each *X* influences *Y*”, “the change in *Y* given a one unit change in *X*”, “how the *X* generated *Y*”) to describe the meaning of the coefficients. Morgan & Winship (2007) and Pearl (2015) give short summaries of this confusion. Spanos (2006a,b) is perhaps the most comprehensive source comparing descriptive vs. explanatory models, parameters, and data generating mechanisms. More generally, Gelman & Hill (2007), Robins *et al.* (2000), and, especially, Shalizi (2017) and Hitchcock (2018) are very accessible accounts of the difference between probabilistic and causal conditioning and statistical vs. causal parameters.

This difference in conditioning between descriptive and explanatory modeling has consequences on inference (Hernán, 2018), including the interpretation of the parameters in a set of nested models in all-subsets regression and on the interpretation of model-averaged coefficients. When the parameters take their meaning from probabilistic conditioning (Equation 4), there is no generating model, simply a statistical model that differs among the nested models. Consequently, the meaning of the parameter *α_j_* differs among the nested models because of different sets of covariates. By contrast, when the parameters take their meaning from causal conditioning (Equation 5), there is a single generating model that applies to all nested models; this is the true, unknown model. Because the parameter for an effector takes its meaning from this unknown generating model, the parameter for *X_j_* has the same meaning in all models and the coefficients from all models are estimating the same parameter. The argument that model-averaged coefficients are meaningless because the coefficients from different models are conditioned on different covariates and, therefore, estimate different parameters, is true *only if* one is descriptive modeling. But, if one is model-averaging the coefficients, or simply, some form of model selection, then the goal of the modeling is probably causal, even if the authors have been inculcated by the norms of their research community that their coefficients have an association interpretation only (Wasserman, 1999; Hernán, 2018). And if the goal is causal (explanatory inference), then coefficients from different models are estimating the same parameter – the casual parameter – and averaged coefficients are meaningful.

## Regression coefficients from different models have the same units

In his influential 2015 criticism of model-averaged regression coefficients, Cade (2015) developed a unique argument against model averaging. Specifically, Cade argued that model averaging is invalid if there is *any* correlation among predictors because model-averaged coefficients “have no defined units in the presence of multicollinearity.” This was not a caution of estimation in the presence of high collinearity (the focus of other collinearity papers) but an argument that all model-averaged coefficients from any observational study are uninterpretable regardless of the magnitude of correlation among the predictors. It is therefore imperative that we understand what Cade means by “no defined units.” This might mean that 1) the units of *b_j.k_* differ among models, or 2) a unit difference in *X_j_* differs among models (Table 1). The first interpretation is simply false; a regression coefficient *b_j.k_* has the units of the simple regression coefficient of *Y* on *X_j_* regardless of the other predictors in the model and there is nothing in the *units* of a regression coefficient that contains information that the value of the coefficient is conditional on some set of covariates. Cade clarifies the second interpretation using the Frisch-Waugh decomposition of **b** = (**X**^T^**X**)^−1^(**X**^T^**y**)

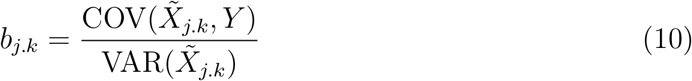

where 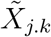 is the component of the variation of *X_j_* not shared with the other *X*, which is simply the vector of residuals of the regression of *X_j_* on the set of covariates *X_k_*. Because the unshared variance 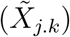 shrinks and swells from model to model, I interpret Cade as stating (Table 1) that the units of *X_j_* itself shrinks and swells from model to model. And, consequently, a unit difference in *X_j_* shrinks and swells from model to model. If it is invalid to average coefficients computed from Equation 10 because of variation in the denominator, then we should not average coefficients from the same model applied to different samples, as these denominators will also differ from sample to sample. But this is moot, because the magnitude of a unit or unit difference is defined by the actual units and not the variance of 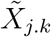. The units of the residuals of weight on height are full kilograms, not partial kilograms. In the owl example, if *ArrivalTime* is measured in hours, a one unit difference is one hour regardless if we are referring to the raw measures or the residuals of *ArrivalTime* on *ParentSex*. One hour (or one kilogram or one degree Celsius) does not shrink or swell among models due to differences in the magnitude of 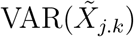.

**Table 1:**
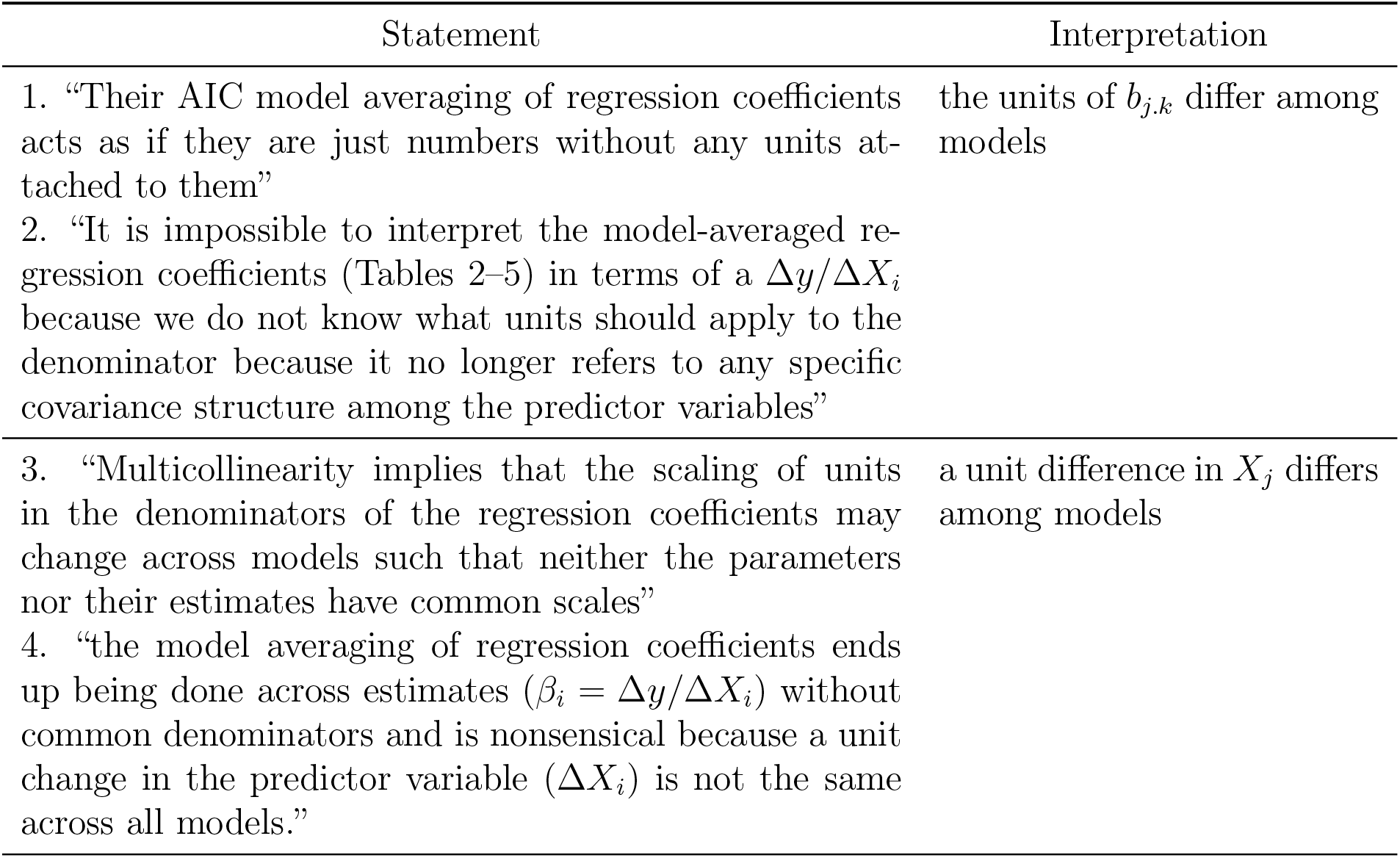
Two interpretation of statements in Cade (2015)

## Conclusion

Statistical models have multiple purposes, including description, prediction, and causal explanation (Mac Nally, 2000; Shmueli, 2010). This variation in purpose is a source of much confusion on the interpretation of regression coefficients both in the primary literature and textbooks because there seems to be little familiarity with the distinction between probabilistic and causal conditioning. In much of the ecology literature, the purpose of the modeling is not explicitly stated and is difficult to infer because the language used is ambiguous. Ecologists often avoid using causal language even if the questions pursued are, ultimately, causal. Larry Wassermen’s comment that “there is always a causal interpretation lurking behind an association” (Wasserman, 1999) is often true in ecology – the owl example above is a case in point. Here, I have focused on statistical modeling for estimating parameters with causal meaning and the “interpretation” of these parameters. Interpretability of model coefficients is not a concern in purely predictive models *used for prediction only* and, consequently, researchers can exploit non-parametric and nonlinear models for improved prediction (Breiman, 2001). By contrast, if we are modeling a system that we hope to manage with manipulations of key variables, or if key variables are changing due to anthropogenic effects (warming, acidification, urbanization), or if we simply want to understand how a system works, then we want our predictive models to estimate parameters with causal meaning. I have argued that, if one is explanatory modeling, a model-averaged coefficient from nested statistical models has a straightforward interpretation as an estimate of the causal effect parameter (Equation 5) from the (generally) unknown true, generating model. While this estimate is conditional on the other covariates in the model, its interpretation or “meaning” is not. Its interpretation takes its meaning from “causal conditioning” and not “probabilistic conditioning.”

Averaging coefficients that potentially estimate different parameters is not unique to model-averaging of coefficients from nested statistical models. Meta-analysis averages effects from different studies and these effects are commonly estimated with different sets of covariates. This averaging is justified because the coefficients are estimating causal parameters that derive their meaning from causal and not probabilistic conditioning. More problematically, meta-analysis averages effects across samples with different distributions of measured confounders. Because of this, Pearl (Pearl, 2012; Pearl & Bareinboim, 2014) criticizes meta-analysis for averaging apples and oranges. But Pearl is not arguing that averaged effects from a meta-analysis fail to have a straightforward interpretation, only that we need more better tools for combining estimates. Regardless, this more insidious issue of meta-analysis is not relevant to model-averaged coefficients of nested statistical models because the same sample is used for all estimates.

Some reviewers of earlier versions of the manuscript have been negative – not because they offered a critique of my argument that coefficients from nested statistical models estimate the same causal effect parameter – but because of 1) issues related to how to model the prior if Bayesian model averaging or how to compute a weight if frequentist model averaging, 2) issues related to reliable estimates of standard errors, and 3) issues with generalized linear models in which a prediction computed as the average of predictions on the response scale is not equal to a back-transformed prediction computed from averaged coefficients on the link scale. These are interesting critiques of model-averaging but not relevant to either the different-parameters criticism of model-averaged coefficients or to my argument developed here. Two other very visceral criticisms on the manuscript have been 1) model-averaging encourages mindless modeling, and 2) model-averaging is a non-starter because it averages over models we know to be wrong (that is, we know they have missing confounders). Again, neither of these critiques addresses what I have written. Nevertheless, both are worthy of a short reply. Mindless modeling is not a property of an estimator but of a researcher, and no strategy is immune to mindless-modeling. Model-averaging is a tool, like penalized regression (including ridge regression), hierarchical regression, Bayesian regression, or structural equation modeling with causal graphs, all of which can and are used quite mindlessly. Using extensive prior knowledge from theoretical and experimental models, a careful researcher will construct a causal model with one or a few, focal predictor variables and all variables that are believed using prior knowledge to be important confounders of the focal effects. The goal is, given this causal model, what can we infer about the effects? The researcher then chooses a tool to estimate these effects.

But why model-average instead of estimating the parameters from what is believed to be the true, generating model, since we know model-averaging averages over models that we know to be wrong? A short answer is: the same reason we use the mid-season batting average of 18 baseball players to predict the end-of-season average of Roberto Clemente and not just Clemente’s mid-season average (Efron & Morris, 1975, 1977). That is, model-averaging is an empirical shrinkage estimator and it is quite easy to show using simulation that under conditions of small to moderate signal-to-noise ratio, model-averaging, which averages over incorrectly specified models, outperforms OLS or ML estimates of the correctly specified model. But again, and it is hard for me to state this more plainly, I am not advocating that model-averaging is the optimal tool, or the best shrinkage estimator, for estimating the effect parameters in explanatory models, only that it is a tool that generates coefficients with straightforward interpretations.

## Acknowledgments

I thank Dr. Brian Inouye, Dr. Brian Cade, and multiple anonymous reviewers for improving the clarity of the manuscript.

## Supplement

The supplement is available at https://doi.org/10.6084/m9.figshare.7967177.v1

