## Supplement 1 for "Model-averaged regression coefficients have a straightforward interpretation using causal conditioning"

### Supplement to: On Model Averaging Regression Coefficients

#### setup

```
library(ggplot2)
library(data.table)
library(mvtnorm)
```

#### “there are, literally, an infinite number of generating models with different combinations of effector (”predictor“) variables that could generate precisely the same probabilistic effect ($b_1$). Here are two”

This header comes from the larger argument that there is not a specific generating model of the data when using probabilistic conditioning because this conditioning makes no assumptions what variables generate the response but only about how the response is related to a specific set of $X$ variables once both are generated and observed.

Here I create fake data generated from different sets of variables but the conditional expectation of the probabilistic effect of $ParentSex$ on the response $NCalls$ conditioned on no other variables is precisely the same.

##### Simplest example – all variables have unit variance and the conditional response is normally distributed.

```
seed_i <- 1
n <- 10^6
rho <- 0.5 # the true correlation between X1 and X2
X <- rmvnorm(n, sigma=matrix(c(1,rho,rho,1), nrow=2))

# Model A - only X1 and U generate the data
# U is the component due to uncorrelated factors that creates "noise"
set.seed(seed_i)
n <- 10^5
beta_1.A <- 0.4
U <- sqrt(1-beta_1.A^2)*rnorm(n)
y1 <- beta_1.A*X[,1] + U
coef(lm(y1 ~ X[,1]))
```

```
##  (Intercept)       X[, 1]
## -0.002056911  0.400308073
```

```
# Model B - X1, X2, and U generate the data
set.seed(seed_i)
n <- 10^5
beta_2.B <- -0.3
beta_1.B <- beta_1.A - rho*beta_2.B
```

```r
sigma_exp <- beta_1.B^2 + beta_2.B^2 + 2*beta_1.A*beta_1.B*rho
U <- sqrt(1-(sigma_exp))*rnorm(n)
y2 <- (X%*%c(beta_1.B, beta_2.B))[,1] + U
coef(lm(y2 ~ X[,1]))
```

```
##  (Intercept)       X[, 1]
## -0.001119979  0.400302096
```

##### More general example – No variables have unit variance and the conditional response is normally distributed.

E($b_1$) for both models is 0.4. The generating effect $\beta_1$ in Model B is 0.55.

```r
n <- 10^6
seed_i <- 1

# create two columns of X that have a correlation of rho
rho <- 0.5
sigma_1 <- 0.5
sigma_2 <- 1.2
sigma.sq_12 <- rho*sigma_1*sigma_2 # cov
Sigma <- matrix(c(sigma_1^2, rho*sigma_1*sigma_2, rho*sigma_1*sigma_2, sigma_2^2), nrow=2)
X <- rmvnorm(n, sigma=Sigma)

beta_0 <- 10 # add an intercept

# model A without X2, again U is the component due to uncorrelated factors, or "noise"
set.seed(seed_i)
beta_1.A <- 2.2
sigma.sq.explained <- (beta_1.A*sigma_1)^2
R2 <- 0.5
sigma.sq.total <- sigma.sq.explained/R2
U <- sqrt(sigma.sq.total - sigma.sq.explained)*rnorm(n)
y1 <- beta_0 + beta_1.A*X[,1] + U
coef(lm(y1 ~ X[,1]))
```

```
## (Intercept)      X[, 1]
##   10.000052    2.200432
```

```r
# model B - with X2
set.seed(seed_i)
beta_2.B <- -0.7
beta_1.B <- beta_1.A - sigma.sq_12/sigma_1^2*beta_2.B
sigma.sq.explained <- (beta_1.B*sigma_1)^2 + (beta_2.B*sigma_2)^2 +
  2*beta_1.B*beta_2.B*rho*sigma_1*sigma_2
U <- sqrt(sigma.sq.total - sigma.sq.explained)*rnorm(n)
y2 <- beta_0 + (X%*%c(beta_1.B, beta_2.B))[,1] + U
# summary(lm(y2 ~ X))$r.squared
coef(lm(y2 ~ X[,1]))
```

```
## (Intercept)      X[, 1]
##   10.000143    2.200426
```

$\hat{b}_1$ for Model A is 2.2004322

$\hat{b}_1$ for Model B is 2.2004259

#### Generalized example – fake owl data

Here the conditional response of generating model is Poisson. The second generating model is constructed to have the same $P(NCalls|ParentSex)$ as the first generating model by

1. setting $\beta_2$ to some value.
2. setting $\beta_1^{(B)} = \beta_1^{(A)} - \frac{\sigma_{12}^2}{\sigma_1^2}\beta_2^{(B)}$.
3. setting $\beta_0^{(B)} = \beta_0^{(B)} - \sigma_{added}$. The subtracted term is the added variance due to the second factor and shifts the intercept on the response scale back to that in model A.

```
seed_i <- sample(1:10^4,1)
n <- 10^6
z_code <- rep(c(0,1),each=n) # the sex "gene" 1=male

# set up mean for females
# exp_beta_0 <- 175 # Ncalls=175, mean calls in females on response scale
# beta_0 <- log(exp_beta_0) # mean calls in log space
beta_0 <- 5.1 # exp(b_0) = 164.02

# set up sex
sex_code <- z_code # i.e. no error mapping gene to sex
sigma.sex_code <- 0.5*sqrt(n*2/(n*2-1)) # sigma(sex_code)
ParentSex <- ifelse(sex_code==0,'F','M')

# set up correlation between arrival time and sex
beta_z <- 4.5 # 4.5 original effect of sex on arrival time
sigma.arrival.unexplained <- 3
ArrivalTime <- beta_z*z_code + rnorm(n*2, sd=sigma.arrival.unexplained) # females centered
# at zero
sigma.arrival_time <- sqrt(beta_z^2*sigma.sex_code^2 + sigma.arrival.unexplained^2)


# expected correlation between sex and arrival time in log space
rho_12 <- beta_z*sigma.sex_code/(sqrt((beta_z*sigma.sex_code)^2 +
                                        sigma.arrival.unexplained^2))
sigma.sq_12 <- rho_12*sigma.sex_code*sigma.arrival_time # covariance
# this generates expected correlations of .6
# cor(ArrivalTime, sex_code)
# cov(ArrivalTime, sex_code)

# these should generate the same coef of sex
# model A - no arrival time
set.seed(seed_i)
beta_1.A <- -0.04 #-0.04
mu.A <- beta_0 + beta_1.A*sex_code # expected log count
count.A <- rpois(n*2, lambda=exp(mu.A))
m1 <- glm(count.A ~ ParentSex,
          family=poisson(link = "log"),
          na.action=na.fail)
coef(summary(m1))
```

```
##                  Estimate   Std. Error    z value Pr(>|z|)
```

```
## (Intercept)  5.09982920 7.808815e-05 65308.6156        0
## ParentSexM  -0.03995618 1.115531e-04  -358.1808        0
```

```
exp(predict(m1, newdata=data.frame(ParentSex=c("F", "M"))))
```

```
##        1        2
## 163.9939 157.5705
```

```
# model B - with arrival time
set.seed(seed_i)
beta_2.B <- 0.05 #.05
beta_1.B <- beta_1.A - sigma.sq_12/sigma.sex_code^2*beta_2.B
# expected added variance

sigma.sq.mu.B <- (beta_1.B*sigma.sex_code)^2 + (beta_2.B*sigma.arrival_time)^2 +
  2*beta_1.B*beta_2.B*sigma.sq_12
sigma.sq.mu.A <- (beta_1.A*sigma.sex_code)^2
sigma.sq.add <- (sigma.sq.mu.B - sigma.sq.mu.A)/2

#beta_0.B <- beta_0 - 0.011 # shifts to same P(NCalls|ParentSex)
beta_0.B <- beta_0 - sigma.sq.add # in response space this is dividing by variance
mu.B <- beta_0.B + beta_1.B*sex_code + beta_2.B*ArrivalTime # expected log count
count.B <- rpois(n*2, lambda=exp(mu.B))
m2 <- glm(count.B ~ ParentSex,
          family=poisson(link = "log"),
          na.action=na.fail)
coef(summary(m2))
```

```
##                Estimate   Std. Error    z value Pr(>|z|)
## (Intercept)  5.09974609 7.809158e-05 65304.6864        0
## ParentSexM  -0.03981013 1.115538e-04  -356.8692        0
```

```
exp(predict(m2, newdata=data.frame(ParentSex=c("F", "M"))))
```

```
##        1        2
## 163.9803 157.5804
```

```
# expected
beta_1.B + sigma.sq_12/sigma.sex_code^2*beta_2.B
```

```
## [1] -0.04
```
